# Chilling injury to algal symbionts induces host starvation and metabolic reorganization in a temperate cnidarian

**DOI:** 10.64898/2026.03.04.709358

**Authors:** Marie Legain, Taissa Lopes Damasceno, Slimane Chaïb, Miriam Reverter, Héloïse Gauthier, Colin Sten Moldenhauer, Paula Isabel Hueso-Jiménez, Suzanne C. Mills, Delphine Raviglione, Nils Rädecker, Nathalie Tapissier-Bontemps, Claudia Pogoreutz

## Abstract

Global temperature anomalies increasingly disrupt the cnidarian-algal symbiosis through a phenomenon termed bleaching. In contrast to heat stress, the mechanisms underlying symbiotic breakdown under cold stress remain largely unknown. Combining physiological and metabolomic measurements, we investigated the response of the photosymbiotic sea anemone holobiont *Aiptasia couchii* to an experiment mimicking a cold spell in the Mediterranean Sea. Within four weeks, we observed the onset of symbiotic breakdown reflected in reduced algal endosymbiont density and chlorophyll *a* content. While photosynthetic efficiency remained largely unaffected, no gross photosynthesis was detectable in cold-stressed anemones and decreases in glycosyldiacylglycerols and fatty acyl glycosides indicated chloroplast lipid remobilization. This breakdown of symbiotic carbon cycling was reflected in increased dipeptide and ceramide levels suggesting anemones catabolized protein reserves and induced pre-apoptotic pathways. Taken together, these responses suggest a decoupling of light and dark reactions of photosynthesis in cold-stressed endosymbionts, resembling chilling injury in higher plants and free-living microalgae. This chilling-induced collapse of symbiotic nutrient cycling eventually leads to host starvation in cold-stressed Cnidaria. Hence, while cold and heat stress may invoke contrasting physiological effects on endosymbionts, our results suggest that both stressors destabilize the symbiosis through similar mechanisms culminating in host starvation.

## Introduction

Coral reefs are among the most biodiverse and productive ecosystems on Earth, supporting an estimated one million species (Knowlton *et al*., 2010). Their ecological success largely depends on the symbiosis between corals and dinoflagellate endosymbionts of the family Symbiodiniaceae (Davy *et al*., 2012). In this mutualistic association, the endosymbionts fix substantial amounts of organic carbon through photosynthesis, and translocate in excess of 40% of their photosynthate, specifically sugars and lipids, to the host, largely meeting the majority of its metabolic demands (Muscatine 1990; Tremblay et al. 2012; Ezzat et al. 2015; Yee et al. 2025). In return, endosymbionts benefit from a stable intracellular environment and access to host-derived metabolic waste products such as inorganic nitrogen and other nutrients, facilitating an efficient nutrient recycling within the holobiont. This tightly integrated metabolic cnidarian-algae symbiosis, however, is highly sensitive to environmental change. Coral bleaching, the breakdown of this relationship with the loss of endosymbionts, is a hallmark of stress most commonly observed under heat stress (Weis, 2008). Heat-driven mass coral bleaching events have increased in frequency and severity since the early 1980s, inevitably driving global coral reef ecosystem loss (Hoegh-Guldberg, 1999). Cnidarian bleaching during heat stress involves a complex cascade of interconnected processes involving impaired photophysiology, oxidative stress, disrupted metabolic exchange, activation of host immune responses, and eventually host starvation. As such, bleaching outcomes are shaped by diverse factors such as host and symbiont functional traits, their respective nutritional status and emerging holobiont properties depending on environmental light, temperature and nutrient regimes (Lesser 1997; Weis 2008; Davy et al. 2012; Rädecker et al. 2021; Pfab et al. 2024). This complex network of interactions limits our mechanistic understanding of the bleaching cascade (Helgoe et al. 2024). However, among these processes, two proposed mechanisms have received particular attention: a) oxidative stress, which involves the overproduction of reactive oxygen species (ROS) by the endosymbionts and/or the host, causing cellular damage to host cells (oxidative theory of bleaching) (Lesser, 1997); and b) the breakdown of mutualistic nutrient cycling between the host and endosymbionts (resource competition hypothesis), which is increasingly recognized as a central mechanism of bleaching (Rädecker *et al*., 2021; Toullec *et al*., 2024; Dubé *et al*., 2025). Specifically, heat stress reduces the photosynthetic capacity of the endosymbionts, which in turn retain photosynthate, effectively reducing carbon translocation to the host. This disruption of symbiotic nutrient cycling ultimately leads to host starvation and, eventually, mortality (Rädecker *et al*., 2021; Toullec *et al*., 2024).

While the processes linking heat stress and symbiosis collapse are fairly well understood, our understanding of the role of other environmental triggers of bleaching and the processes involved remains limited. Critically, only few studies have focused on bleaching caused by low temperatures to date (Steen & Muscatine, 1987; Hoegh-Guldberg *et al*., 2005; Lirman *et al*., 2011; Pontasch *et al*., 2017; Marangoni *et al*., 2021; Rich *et al*., 2022; El-Khaled *et al*., 2025). Marine cold spells remain an underinvestigated, yet geographically wide-spread phenomenon which has been reported to affect reef-building corals from the Caribbean to the Indo-Pacific, including the Central Red Sea and the Mediterranean Sea (Hoegh-Guldberg et al. 2005; Kemp et al. 2011; Serrano et al. 2017; Rich et al. 2022). In general, these extreme cold events can be of comparable magnitude as major marine heatwaves (Lentini et al. 2001; Florenchie et al. 2004) and as such are ecologically important phenomena which can shift the distribution of species, alter benthic community composition (Schlegel et al. 2021), and cause bleaching in photosymbiotic Cnidaria (Saxby et al. 2003; Hoegh-Guldberg and Fine 2004). Notably, in contrast to marine heatwaves, marine cold spells currently experience a decrease in frequency, severity, and duration, with the exception of the Southern Ocean (Schlegel et al. 2021; Wang et al. 2022). However, as some photosymbiotic scleractinian corals and sea anemones exhibit trends towards poleward migration (Yamano et al. 2011; Malcolm and Scott 2017; Landry Yuan et al. 2023), it is important to understand their lower temperature limits, including their bleaching response to (severe) cold stress.

While cold-induced bleaching in photosymbiotic Cnidaria has received less attention, it shares remarkable similarities with heat-induced bleaching. Both can result in decreased photosynthetic efficiency (Saxby *et al*., 2003), affect endosymbiont densities (Saxby *et al*., 2003; Rodríguez-Troncoso *et al*., 2010; Roth *et al*., 2012; Nielsen *et al*., 2020), and reduce growth (Roth *et al*., 2012) and respiration rates (Higuchi *et al*., 2015). In contrast to heat-induced bleaching though, the metabolic and molecular processes underpinning cold stress in the cnidarian holobiont remain largely unknown (Saxby *et al*., 2003; Roth *et al*., 2012). Importantly, heat stress induces notable shifts in holobiont metabolite profiles, specifically increased abundances of stress-related metabolites (phospholipids, steroids, organic acids) and declines in compounds linked to steady-state physiological activity (signalling compounds, structural lipids, antioxidants) (Hillyer *et al*., 2016; Farag *et al*., 2024, 2025). To date, no studies integrating changes in metabolite profiles with physiological responses under cold stress are available for photosymbiotic cnidarians.

The strong seasonality of the Mediterranean Sea constitutes a unique natural laboratory to study the effects of thermal stress on photosymbiotic Cnidaria (Coll *et al*., 2010). With annual temperature amplitudes exceeding 15°C in many places (Rodolfo-Metalpa *et al*., 2006), recurrent bleaching episodes occur not only during heat waves in the summer but also as a result of cold stress during winter months (Serrano *et al*., 2017). Cold spells -very cold winters and frost events - occur about once every decade in the Mediterranean (Larcher, 2000). In general, both chronic and abrupt cold-water events have detrimental effects on the physiology of photoautotrophic organisms (Manasa *et al*., 2022) and marine life, and have the capacity to cause bleaching and mortality (Green *et al*. 2019; Riegl *et al*., 2019). Ultimately, recurring alternating heat waves and cold spells could therefore pose a serious threat to vulnerable benthic Mediterranean ecosystems.

Photosymbiotic sea anemones are powerful models for functional studies on coral bleaching (Lehnert *et al*., 2012), as they exhibit similar physiological responses (Grajales & Rodríguez, 2014), but can be more easily maintained and experimentally manipulated in the lab than most corals (Rädecker *et al*., 2018). We here set out to evaluate the physiological responses of *Aiptasia couchii* holobionts from the Western Mediterranean to cold stress by combining physiological responses (oxygen fluxes, symbiont density, maximum quantum yield of the algal photosystem II) with bulk metabolomic (i.e., holobiont: without separating the anemone from its endosymbionts) analysis. This integrated approach permitted the assessment of cold stress effects on the photosymbiotic holobiont, specifically on symbiotic nutrient cycling. We hypothesized that cold stress causes bleaching in the *A. couchii* holobiont by disrupting symbiotic nutrient cycling, culminating in host starvation.

## Material and methods

### Model system

Trumpet anemones of the family Aiptasiidae constitute a monophyletic group (Grajales & Rodríguez, 2016). We here employed *A. couchii*, a common species which forms extensive clonal patches in the shallow rocky intertidal of the Mediterranean Sea (Arossa *et al*., 2021) and harbours an uncharacterized endosymbiotic member of the Symbiodiniaceae: *Philozoon* sp. (’temperate *Symbiodinium*’) (LaJeunesse *et al*., 2022).

### Study area, animal collection and acclimation

Surface water temperatures in the Western Mediterranean Sea typically range from approximately 10°C during the coldest months to above 28°C during peak summer, particularly in shallow coastal areas where thermal stratification is most pronounced (Shaltout & Omstedt, 2014). To determine the natural thermal environment *in situ* during the coldest time of the year, a HOBO temperature logger was deployed near the collection site between Collioure and the Plage de l’Ouille (’Pont Cassé’, France; 42°31’57.1"N 3°04’49.8"E) (**Supp. Figure S1A**). Water temperatures were recorded for several weeks (27 January - 10 March 2025) prior to the sampling campaign to document the minimum temperatures. The lowest temperature measured during this period was around 11°C (**Supp. Figure S1C**).

*Aiptasia couchii* were collected using hammer and chisel from the rocky shore in March 2025 (**Supp. Figure S1D**). Collection permits were issued by the Direction Interrégionale de la Mer Méditerranée (permit decision number 205). After transfer to the lab, sea anemones were maintained in transparent plastic containers (2.25 L capacity each, Rotho®, Switzerland) each filled 0.22 µm filtered artificial seawater (Tropic Marin Pro-Reef; salinity 35 psu) and maintained at 17°C to gently acclimate for two weeks. This husbandry and control temperature was chosen based on the annual mean sea surface temperature for Collioure for the period 2015-2025 (Copernicus Marine Services; **Supp. Figure S1B**; **Supp.Table S1**). Sea anemones were maintained in temperature-and light-controlled incubators (FitoClima 600 PDH Bio 2, Aralab®, Portugal) under a 12:12 h light-dark cycle, with an irradiance of 42 μmol photons m⁻² s⁻¹ measured using a quantum sensor (Apogee Quantum Flux, USA). During the acclimation period, individuals were fed once per week ad libitum in the evening with freshly hatched *Artemia salina* nauplii. The seawater in each container was replaced and biofilms on all surfaces removed with laboratory-grade cotton swabs each morning following feeding.

### Experimental design for cold stress experiment

A total of 66 *A. couchii* (range of oral disc diameters 5-20 mm) were used to determine the effects of cold stress on holobiont physiology and metabolomic response. At the end of acclimation, the selected anemones were randomly assigned to two conditions: control (n = 33) and cold stress (n = 33). For each condition, anemones were distributed across three replicate clear plastic containers (400 ml capacity each; n = 11 per condition) filled with filtered (0.22 µm) artificial seawater (salinity 35 psu).

The control anemones were maintained at 42 μmol photons m^-2^s^-1^ with a 12:12h light-dark cycle, at 17°C. The anemones subjected to cold stress were maintained under the same light conditions. Following a test experiment to determine a non-lethal low temperature stress that caused a clear phenotype in exposed anemones (Supplementary information for details; **Supp. Figures S2A, S3**), a cold stress was applied through gradual, non-linear downramping of incubator temperatures (**Supp. Figure S2B**) over a period of four weeks. Anemones were not fed during the experiment. Twice a week, containers were cleaned with laboratory-grade cotton swabs and replenished with filtered seawater. Throughout the duration of the experiment, phenotypic observations (onset of bleaching, tentacle retraction, mortality) were conducted weekly around 6 pm.

### Photosynthetic performance of holobionts

To assess effects of temperature treatments on algal photophysiology, we performed measurements of the maximum quantum yield (Fv/Fm) of photosystem (PS) II of endosymbiont communities *in hospite*, a rapid, non-invasive parameter indicative of PSII health (Strasser et al., 2020). The Fv/Fm was measured via PAM fluorometry (Pulse-Amplitude Modulation) with a JUNIOR-PAM (red version; Walz®, Germany) once a week in the dark (one hour before the lights turned on in the morning), just prior to the next decrease in temperature in the cold-stress treatment (**Figure 2**). This design allowed for consistent weekly monitoring of the status of PSII health of endosymbionts throughout the experiment (n = 9 anemones per condition).

### Photosynthesis and respiration rates of holobionts

At the end of the experiment, oxygen (O_2_) flux measurements of control and cold-stressed anemones were used to estimate net photosynthesis and respiration rates in closed light and dark incubations, respectively. Individual anemones were transferred into custom-made acrylic incubation chambers (15 ml) and settled onto a stand, and a magnetic stirring bar was placed underneath. Incubation chambers were filled with artificial seawater, transferred to treatment incubators (17°C or 6°C, respectively), and placed on magnetic stirring plates (225 rpm, Cimare i Poly 15 and Multipoint Stirrers, Thermo Fisher Scientific, USA) for constant homogenization of the water column. O_2_ concentrations were recorded every second for approximately 1 hour at a constant light level of 42 μmol photons m⁻² s⁻¹ (mean irradiance during the 12-hour light phase), followed by approximately 1 hour in the dark, using adhesive O_2_ sensor spots in the chamber connected to FireSting O_2_ optical oxygen meters (PyroScience, Germany).

To obtain O₂ fluxes, local linear regressions were performed using the *LoLinR* R package (Olito *et al*., 2017), which objectively identifies the optimal linear portion of time-series data for estimating O₂ fluxes during light and dark incubations. Slopes of linear regressions represented the rate of change in O_2_ concentrations over time, corresponding to net photosynthesis (P_net_; derived during light incubations) and dark respiration (R; dark incubations). These rates were then corrected by subtracting values obtained from blanks (i.e., incubation chambers filled with seawater only), to account for pelagic background metabolism. Corrected fluxes were normalized by chamber volume and host tissue protein content. Gross photosynthesis (P_gross_) rates, representing the total amount of O_2_ produced during photosynthesis, were calculated as the sum of the P_net_ rates and the absolute value of the dark respiration rate of individual anemone holobionts.

### Preparation of anemone samples for physiology measurements

Six anemones per condition were used to assess host protein content, endosymbiont density, and chlorophyll *a* (chl *a*) content. Excess water and mucus were carefully removed, and anemones placed in a pre-weighed 5 ml Eppendorf tube and weighed using a precision balance. Cold sterile 2 × PBS buffer was added by taking into account anemone wet weight and aiming for a final volume of 1.5 mL: the volume of PBS buffer added was 1.5 mL from which the anemone volume (in microliters), estimated from the anemone wet weight and assuming a density of 1 g.mL^−1^, was subtracted. The anemones were then homogenized on ice in the 2 × PBS buffer using a sterile pestle and a hand-held Pellet Pestle Cordless Motor (DWK Life Sciences, USA) until the tissue slurry was homogeneous. To separate the host and endosymbiont fractions, the homogenate was centrifuged (3,000 × g, 4°C, 5 minutes) (Allegra X-30R Centrifuge, Beckman Coulter®, USA). The supernatants containing the host fraction were split into two aliquots, transferred into 1.5 ml cryogenic tubes, immediately snap -frozen in liquid nitrogen, and stored at −80°C until further processing. Endosymbiont pellets were purified by two steps of centrifugation (3,000 × g, 4°C, 5 minutes each) and resuspension of pellets in 1 mL of 2 × PBS buffer. The resuspended cells were separated into two fractions for chl *a* content measurement and endosymbiont density quantification. These samples were stored at −20°C until further analyses.

### Endosymbiont density in host tissues

To assess the density of endosymbionts within host tissues, frozen endosymbiont pellets were gently thawed on ice, resuspended by vortexing, and centrifuged (3,000 × g, 4 minutes). The supernatant was discarded, and pellets were incubated in 1 mL of 1 M NaOH at room temperature to dissolve remaining host tissue debris to increase accuracy of endosymbiont counts (Zamoum & Furla, 2012). Samples were resuspended by vigorous shaking by hand every 15 minutes for one hour, and then centrifuged again (3,000 × g, 4 minutes). After discarding the supernatant, the endosymbiont pellet was washed twice in 1 mL of 2 × PBS buffer. Finally, the pellet was resuspended in 2 × PBS and stored at 4°C until quantitation. Cell counts of four technical replicates were performed with an improved Neubauer chamber using an optical microscope (Leica, Wetzlar®, Germany). Averaged endosymbiont counts for each anemone were normalized to host protein content to obtain cell densities per unit of host biomass.

### Endosymbiont chlorophyll *a* content

As chl *a* is the primary pigment involved in photosynthesis, its concentration serves as a reliable proxy for the photosynthetic potential of endosymbiont cells. The chl *a* content of endosymbiont cells was determined from an aliquot of 700 μL of the cell suspension. Samples were gently thawed on ice, resuspended by vortexing and then centrifuged (3,000 × g, 4 minutes). The supernatant was discarded and the resulting pellet was resuspended in 500 μl of ice-cold ethanol (96%). The samples were incubated in the dark at 4°C under constant rotation overnight for chlorophyll extraction. After centrifugation (3,000 × g, 4°C, 5 minutes), chlorophyll extracts were transferred into a flat bottom 96-well plate for absorbance measures. Ethanol (96%) was used as a blank. Measurements were performed in duplicate. The absorbance of each sample was measured at 630, 664, and 750 nm with a plate reader (CLARIOstar Plus, BMG LABTECH®, Germany), and the chl *a* content was calculated as follows, taking into account the turbidity correction (Jeffrey & Humphrey, 1975):

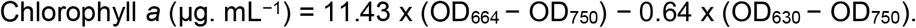

Finally, chl *a* content was standardized to the corresponding host protein content of each individual sample.

### Total host protein content

The tissue protein content of anemones was assessed using the Quick Start Bradford Protein Assay Kit (Bio-Rad®, USA) on one of the frozen aliquots of the endosymbiont-free host supernatant, following the manufacturer’s instructions for the standard microplate protocol. Technical 5 μl triplicates of each sample were measured in a flat-bottom 96-well plate. Protein content was assessed using a plate reader (CLARIOstar Plus, LABTECH, Germany) by measuring the absorbance at 480 nm. The absorbance measurements were calibrated against a serial dilution of Bovine Serum Albumin (BSA) protein standard. The host protein content was used to normalize physiological response parameters.

### Quantitation of superoxide dismutase activity in host tissues

The second aliquot of endosymbiont-free host supernatants was gently thawed on ice for measurements of the antioxidant superoxide dismutase (SOD) activity using the Total Superoxide Dismutase (T-SOD) Activity Assay Kit (WST-1 Method) (E-BC-K020-M; Elabscience, USA). Enzyme solutions were prepared fresh just before the assay. 20 μl aliquots of host supernatant were used for the assay for each biological replicate in a black flat-bottom well plate according to manufacturer’s instructions. SOD activity was then assessed alongside appropriate blanks using a plate reader (CLARIOstar Plus, LABTECH, Germany) by measuring absorbance at 450 nm. Obtained values were then used to calculate SOD activity using the appropriate formula for tissue and cell samples as provided by the kit manufacturer: SOD activity (U/mgprot) = *i* ÷ 50% × *V*1/*V*2 × *f* ÷ *Cpr*

Where i is the inhibition ratio of SOD (%); V1 the total volume of reaction, 240 μL; V2: The volume of sample added to the reaction, 20 μL, f: Dilution factor of sample before test; Cpr: Concentration of protein in sample, gprot/L.

### Untargeted metabolomics of sea anemone holobionts

#### Metabolite extraction and sample preparation

Ten *A. couchii* per condition were used to assess the impact of cold stress on metabolite profiles. Each anemone was snap-frozen in liquid nitrogen immediately after collection and then freeze-dried overnight. Freeze-dried anemones were individually ground into powder using a bead beater (Tissuelyser II, QIAGEN®, Germany) under liquid nitrogen and stored at−80°C until extraction. Due to the low biomass available and the marine origin of the samples, we prioritized a single standardized extraction using a simple and robust approach which maximizes reproducibility, which retains and separates polar and semi-polar metabolites, which minimizes the co-extraction of salts that might cause ionization-related issues during analysis, and which is suitable for Cnidaria (Farag et al. 2024): For extraction, 10 mg of powdered freeze-dried anemone per biological sample were homogenized in 1.0 mL of MeOH (HPLC-MS grade) containing 0.1 μg.ml^-1^ umbelliferone (≥ 98% purity; Sigma-Aldrich, Germany) as internal standard by ultrasonication at room temperature (2 minutes). After homogenization, the samples were shaken for 15 minutes with an automatic shaker (1.500 rpm, Multi Reax Orbital Shaker, Heidolph, Germany) and then centrifuged (4,400 rpm, 4°C, 10 minutes) to remove cellular debris (Farag et al. 2024). For LC-MS analysis, 300 μL aliquots of the supernatant were 1:10 diluted with MeOH (HPLC-MS grade) and filtered (PTFE, 13 mm, 0.22 μm, (J.T.Baker, Avantor, France). Quality control (QC) was implemented by pooling equal volumes of each sample. Two extraction blanks were prepared in parallel with samples to detect potential contaminants. Additional QC samples (condQC) were prepared for each condition by pooling an equal volume of respective samples. All samples were immediately analyzed.

#### UHPLC-HRMS profiling

Sample injection order was randomized, with QC samples injected every five samples to monitor system performance and reproducibility. MeOH blanks, extraction blanks and QC samples were injected at the beginning and at the end of the analytical sequence to ensure column conditioning and system suitability. CondQC samples were analyzed at the very end of the analytical sequence to perform MS/MS analysis in negative and positive ionisation modes.

Following extraction, the samples were injected into a Vanquish UHPLC, a high-performance liquid chromatography (UHPLC) system (Thermo Fisher Scientific, USA), coupled with a QExactive plus Orbitrap mass spectrometer (Thermo Fisher Scientific, USA). Chromatographic separation was performed on a Phenomenex Luna Omega Polar C18 column (1.6 μm, 100 Å, 100 × 2.1 mm) at 40°C. Gradient solvents were composed of water with 0.1% formic acid (A; 99% RS, Carlo Erba, Val de Reuil, France) and acetonitrile (LC-MS grade, Carlo Erba, Val de Reuil, France) with 0.1% formic acid (B) delivered at a flow rate of 0.4 mL/min. The elution gradient was programmed as follows: 5% of B for 2 min, linear increase from 5% to 100% of B in 13 min, 100% (B) for 4 min, from 100% to 5% (B) in 1 min and, re-equilibration at 5% (B) for 5 min (injection volumes 2 μL). Full scan mass spectra were acquired both in positive and negative modes from 1 to 19 min. Mass spectrometer settings were as follows: sheath, auxiliary and sweep gas 45, 15 and 0 AU, respectively; spray voltage, 3,200 V in both positive and negative modes; capillary temperature, 320°C; ESI probe heater temperature, 250°C and S-lens RF level, 60. Full scan mass spectra (MS1) were acquired in both positive (UHPLC-ESI(+)-MS/MS) and negative (UHPLC-ESI(-)-MS/MS) ionization modes with a full scan MS window of 120–1500 m/z and a resolution of 35,000. The maximum Injection Time (IT) was set to 100 ms and automatic gain control (AGC) target set at 1E6. For metabolite annotation and molecular network analysis, MS/MS acquisition consisted of one full scan mass spectrum and 5 data-dependent MS/MS scans. For the Full MS, the resolution was 70,000, Automatic Gain Control (AGC) was 3E6, the maximum IT was 100 ms and a scan mass range of 120-1,500 m/z was used. For the dd-MS^2^/dd-SIM experiments, the resolution was 17,500, AGC target was 1E5, maximum IT was 50 ms and for each MS/MS scans, top 5 most intense ions taking into account an isolation window of 2 m/z and a fixed first mass of 50.0 m/z were fragmented. Three Normalized Collision Energy™ (NCE) values (25 eV, 35 eV, 45 eV) were applied to obtain a combination of 3 fragmentation spectra. Dynamic exclusion was set to 5 s and isotope peaks were excluded. The data were evaluated using XCalibur software version 4.5 (Thermo Fisher Scientific, USA).

#### UHPLC-HRMS data processing and annotation

Raw UHPLC-HRMS data obtained in positive and negative ionization modes were converted into mzXML files in centroid mode with MS convert software (from Proteowizard suite, version 3.0.2, Palo Alto, CA, USA). MzXML files were directly processed using MZmine 4 (7.9 version) (Schmid *et al*., 2023). Data obtained in UHPLC-ESI(+)-MS/MS (positive) and UHPLC-ESI(-)-MS/MS (negative) ionization modes were processed separately. Initially, positive and negative mode data were analysed using exclusively MS1-level data to conduct biostatistical analysis and biomarker discovery. Subsequently, only condQC were subjected to tandem mass spectrometry (MS/MS) fragmentation to generate MS/MS data. These fragmentation data were used to perform spectral similarity networking and metabolite annotation.

Masses were detected in centroid mode at a minimal intensity threshold (MS1:1.0 × 10⁴; MS2: 1.0 × 10²). An Automated Data Analysis Pipeline (ADAP) module was employed to construct chromatograms with the following parameters: minimum group size of four scans, minimum peak height of 1.05, and m/z tolerance of 20 ppm. For chromatogram deconvolution using a local minimum feature resolver, the following settings were applied: minimum absolute height of 1.0 × 10³, minimum data points of 3 scans, peak duration range of 0.01–0.5 min, and minimum retention time (RT) search range of 0.05 min. Isotopic peaks were removed using the isotopic peak grouper with the following parameters: m/z tolerance of 10 ppm, RT tolerance of 0.1 min, maximum charge of 2, and selection restricted to the most intense monoisotopic ions. Chromatograms were subsequently aligned with m/z and RT tolerances of 10 ppm and 0.1 min, respectively, with equal weighting (1.0) assigned to both m/z and RT dimensions. Isotope pattern comparison was performed using the following thresholds: isotope m/z tolerance of 10 ppm, minimum intensity of 10, and minimum pattern score of 90%. The resulting aligned chromatogram list was filtered to remove duplicate peaks using m/z tolerance of 5 ppm and RT tolerance of 0.1 min. A selection criterion was applied to retain only isotopic patterns containing at least two peaks occurring in at least two samples. Gap filling was conducted using the Feature Finder algorithm with an intensity tolerance of 10%, m/z tolerance of 10 ppm, and RT tolerance of 0.1 min, followed by RT correction. Blank sample filtering was performed to remove all features detected in extraction blank samples, applying a minimum detection threshold of two blanks and quantification based on peak area integration. MS1 peak intensities (i.e., peak areas) from each of the two datasets were exported to .csv format files for statistical analysis. For the second dataset containing MS/MS fragmentation data, a table comprising MS1 peak intensities of features with corresponding MS2 data and an .mgf file containing consensus MS/MS spectra were exported for spectral similarity network construction and metabolite annotation via the Global Natural Products Social molecular networking web-platform (GNPS) (Wang *et al*., 2016) and SIRIUS (Dührkop *et al*., 2019).

Molecular formulas of the most significantly differentially abundant features were calculated with Sirius 5.6.2, using Orbitrap base parameters (mass tolerance: 5 ppm; isotopic ratio tolerance: 20%) (Dührkop *et al*., 2019). Compound class annotation was performed using CANOPUS integrated in SIRIUS and the ClassyFire chemical ontology (Djoumbou Feunang *et al*., 2016), and only those with scores above 0.5 were retained. CSI-Finger ID was used to obtain compound structure IDs, which were individually evaluated using the CSI-Finger ID scores and Tanimoto fragmentation coefficients.

### Statistical analyses

#### Physiological data analysis

Statistical significance between controls and cold-stressed holobionts was assessed using a t-test, provided that the assumptions of normality and homogeneity of variances (verified by the Shapiro-Wilk and Levene tests, respectively) were met. When more than two groups were compared, ANOVAs were performed using the “car” package (Fox & Weisberg, 2018), under the same assumptions. When these conditions were met, Tukey’s post hoc test was applied; otherwise, a non-parametric Kruskal-Wallis test was used, followed by Dunn or Conover’s post hoc test for multiple comparisons. Outliers were identified for all physiological parameters and metabolite profiles, and removed for SOD activity using the InterQuartile Range (IQR) method (Vinutha et al., 2018). All statistical analyses were carried out using R Studio (R version 4.3.1) (R Core Team, 2024). The significance level was set to 95% (α = 0.05). All data are expressed as mean ± standard error.

#### Statistical analysis of metabolite data

Data matrices generated from both ionization modes were exported on MetaboAnalyst 6.0 (Ewald *et al*., 2024) for statistical treatment. An initial filtering step was performed using the low-repeatability filter module with a threshold of 30% to remove features with poor reproducibility across QC samples. Low variance filters were applied using Median Absolute Deviation (40% for positive dataset and 20% for negative dataset). Low-abundance filtering (threshold: 10%) was applied using the Median intensity value. Missing value imputation was conducted using the median estimation method from the Univariate Statistical Methods module. Finally, data normalization was applied in two sequential steps: auto-scaling (mean-centering and unit variance scaling) followed by sample-wise median normalization to account for systematic variations in peak intensities across samples. The data matrices for negative and positive ionization modes included 1778 and 2212 features, respectively. Statistical significance of differences in abundances of individual metabolites between control and cold-stressed anemones was assessed using univariate t-tests using the Vulcano analysis function. Metabolites with an FDR-adjusted *p* value < 0.05 and a Fold Change (FC) threshold of 2 were considered significantly increased or decreased in abundance under cold stress conditions. Unsupervised principal component analysis (PCA) in addition to supervised orthogonal projection to least-squares discriminant analysis (OPLS-DA) was employed (Worley & Powers, 2016). OPLS-DA maximizes the separation between the groups, highlighting the importance of each variable based on estimation of the Variable Importance in Projection (VIP) scores. Permutation tests and a double cross-validation test (2CV) were performed to validate the OPLS-DA model. All VIP features with a score above 1.6 were selected for further analysis and manually checked with the MZmine software. An ion extraction at the defined retention time was performed, taking scan number and its isotopic mass into account. VIP corresponding to ions that were not fragmented in MS/MS were discarded. VIP with low isotopic pattern quality were removed and for those with identical exact masses (i.e., isomers), only the isomers with the highest intensity were retained. A Wilcoxon rank sum test was performed to test the significance of the differences in intensity of VIP features between the control and cold-stressed conditions using RStudio environment v4.0.4 (R Development Core Team, 2008).

#### Mass spectral similarity network

The molecular networks from data from both positive and negative ionization mode were generated by converting mzXML files obtained from condQC analysis, using the FBMN online workflow on the GNPS (Wang *et al*., 2016). MS/MS spectra were filtered by choosing the top 6 peaks in the +/−50 Da window throughout the spectrum. For some analysis, data were clustered with MSCluster with a parent mass tolerance of 0.02 Da and a MS/MS fragment ion tolerance of 0.02 Da to generate consensus spectra (Frank *et al*., 2008). Edges between two nodes were kept in the network when each of the nodes appeared in each other’s respective top 10 most similar nodes. The library spectra were filtered in the same manner. All matches retained between network and library spectra were required to show a cosine score above 0.7 and at least six matched peaks. The generated similarity networks were compared to databases as implemented in GNPS (CASMI Spectral Library, GNPS library). Spectral similarity networks generated with GNPS were imported into Cytoscape v3.8.2 for processing (Shannon *et al*., 2003). First, the spectral similarity networks including VIPs metabolites were isolated for each ionization mode. For each mode, metadata were merged and harmonized between the networks to obtain a unique network of similarity. For this, each MS and MS/MS of each ion from GNPS networks were compared pairwise for validation. The connections to the other nodes were harmonized into a mean cosine score derived from the values obtained with GNPS. For this, ions potentially belonging to background noise or being artifacts were ruled out. The Allegro Layout representation mode was used to organize the spatial arrangement of the arrays in Cytoscape (Shannon *et al*., 2003). Global molecular networks obtained from positive and negative modes were scrutinized to highlight clusters grouping chemical classes of metabolites that differ significantly between control and cold-stressed conditions. Mean of summed relative intensities (peak areas) in all the samples were re-assigned to VIPs highlighted in the subnetworks of interest according to their ClassyFire chemical ontology (Djoumbou Feunang *et al*., 2016) from CANOPUS (Sirius) output (Dührkop *et al*., 2021).

## RESULTS

### Holobiont phenotype and symbiont density

Cold-stress had a pronounced effect on *A. couchii* phenotypes on day 28 of the experiment: while control individuals appeared visibly healthy with expanded tentacles in which clusters of endosymbionts were visible (**Figure 1 A,B**), cold-stressed individuals (**Figure 1 C,D**) exhibited retracted tentacles with partial whitening of the tips potentially indicating a localized loss of endosymbionts.

**Figure 1:**
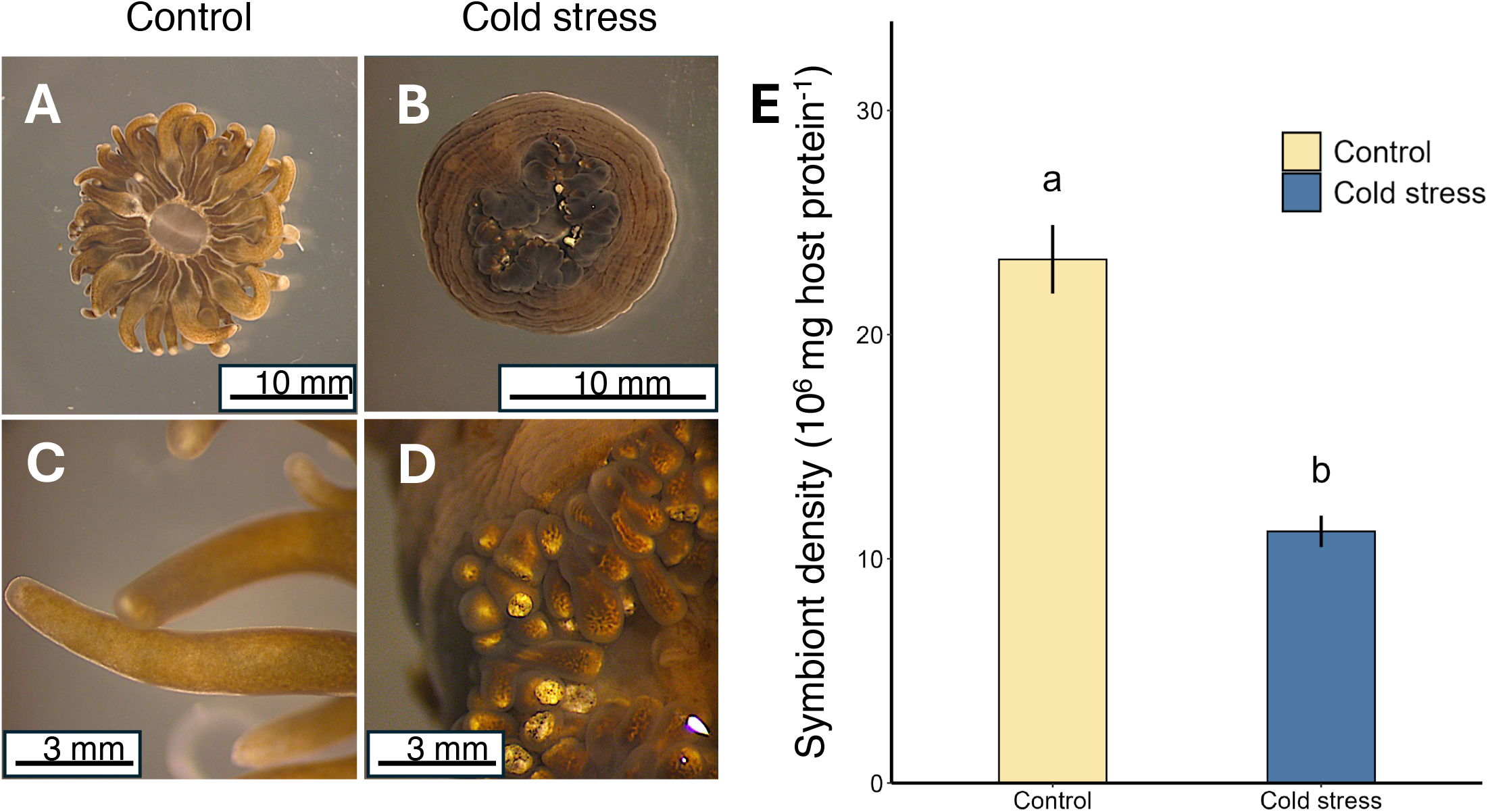
Phenotype and algal symbiont densities of control and cold-stressed sea anemones *A. couchii*. **A,C:** overview shots of entire individuals (top row) and close-ups (bottom row) of tentacles of *A. couchii* holobionts (**B,D**) at the end of the cold-stress experiment (4 weeks): Comparison of non-stressed control (**A, B**, 17°C) and cold-stressed individuals (**C, D**, 6°C). The images were captured using a stereomicroscope. **E**: Symbiont density per mg of host tissue protein in *A. couchii* sea anemone holobionts under two experimental conditions: control and cold stress (n = 6 per condition). Different letters above bars indicate statistically significant differences between groups (*p* < 0.05). Data are presented as mean ± standard error.

In line with this, the density of symbionts per unit of host tissue protein decreased significantly in cold stressed holobionts (11.2 ± 3.44 x 10⁶ symbiotic cells·mg⁻¹ host tissue protein) compared to controls (23.2 ± 7.48 x 10⁶ symbiotic cells·mg⁻¹ host tissue protein) (H = 35.2, p < 0.001, Kruskal-Wallis test), representing a decrease of approximately 52% (**Figure 1E; <u>Supp. Table S2</u>**).

Further, significant mucus production was observed during sampling in the control animals, but not in the cold-stressed animals (*personal observation ML, CP*). No mortality was observed in either condition.

### Chlorophyll *a* content and photosynthetic efficiency of algal endosymbionts

Following trends in symbiont densities, chl *a* content varied significantly between treatments. Specifically, chl *a* content per mg of host tissue protein was significantly lower under cold-stress (2.21 ± 0.29 µg.mg⁻¹ host tissue protein) compared to the control (3.46 ± 1.49 µg.mg⁻¹ host tissue protein), corresponding to a 36% decrease (Z = −2.4707, *p* = 0.011, Fisher-Pitman Permutation Test) (**Figure 2B**). In contrast, chl *a* content per algal cell was significantly higher in the cold stress treatment (2.06 × 10⁻⁷ ± 7.37 × 10⁻⁸ µg.symbiont cell⁻¹) compared to the control (1.60 × 10⁻⁷ ± 7.64 × 10⁻⁸ µg.symbiont cell⁻¹), representing an increase of approximately 29% (H = 4.56, p = 0.033, Kruskal-Wallis test) (**Supp. Figure S4A**; **<u>Supp. Table</u> <u>S2</u>**).

**Figure 2:**
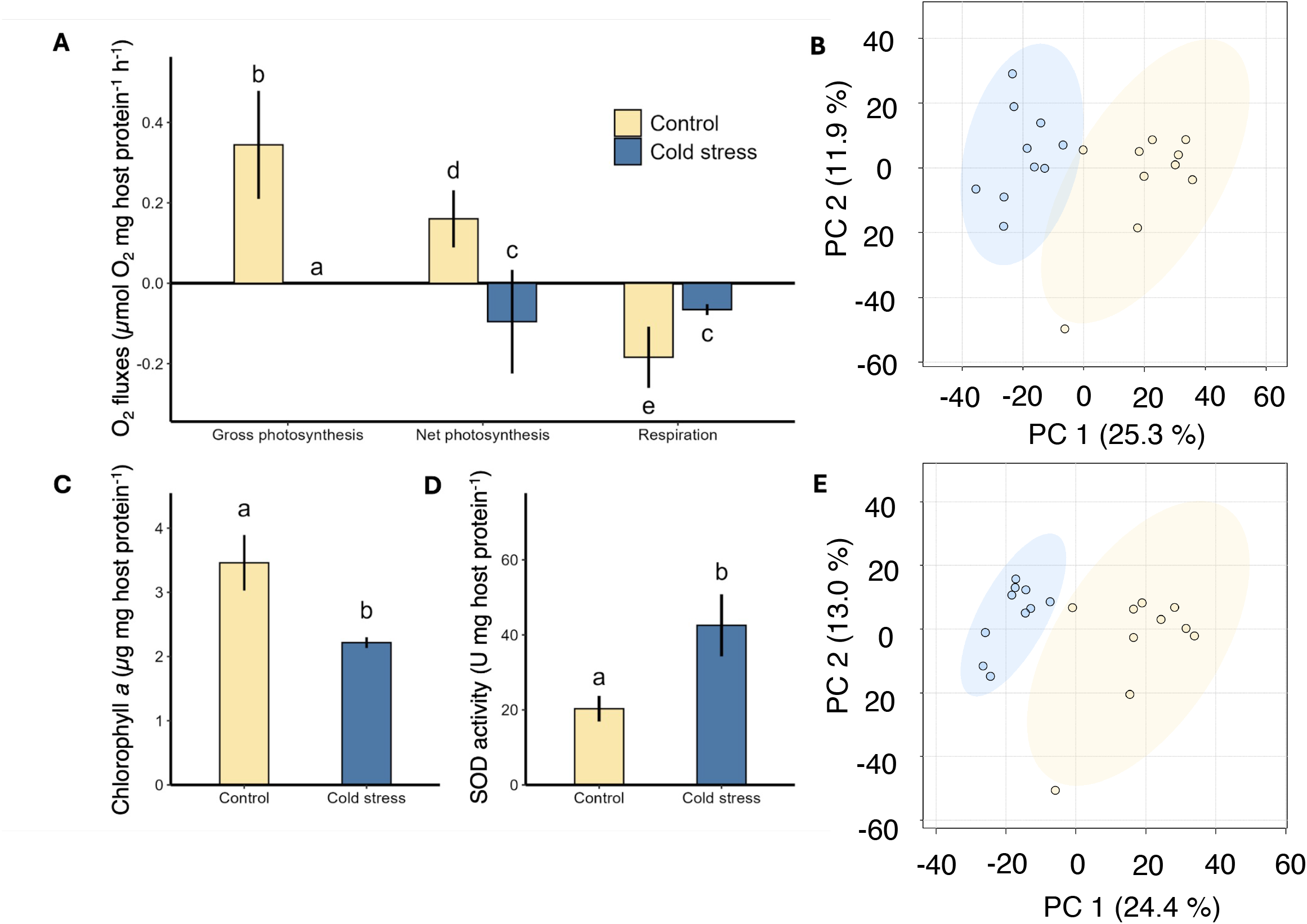
Physiological responses, antioxidant activity, and changes in metabolite profiles in *A. couchii* following four weeks of non-linear cold stress. A: Oxygen (O₂) fluxes related to gross photosynthesis (left), net photosynthesis (center) and dark respiration (right) in control (yellow) and cold-stressed (blue) *A. couchii* (in µmol O₂·mg host tissue protein⁻¹·h⁻¹; n = 6 per condition). **B:** Principal component analysis (PCA) of metabolite profiles of anemone holobionts in positive ionization mode (n = 10 per condition). **C:** Chlorophyll *a* content of algal symbionts normalized to host tissue protein content. **D:** Superoxide dismutase (SOD) activity of *A. couchii* host tissue homogenates normalized to host tissue protein content. **E:** PCA of metabolite profiles of anemone holobionts (i.e., host tissue and algae) in negative ionization mode (n = 10 per condition). Letters indicate statistically significant differences between groups (*p* < 0.05). Data are presented as mean ± standard error.

Despite the clear host phenotype and algal symbiont densities, Fv/Fm remained fairly stable across the experimental groups (**Supp. Figure S4B**; **<u>Supp. Table S3</u>**), ranging from 0.592 ± 0.079 to 0.635 ± 0.038. No significant effects of condition (control vs. cold stress; F = 1.39, *p* = 0.241), temperature (from 17°C to 6°C; F = 1.17, p = 0.328), or their interaction (F = 0.258, p = 0.904, Type II ANOVA, multiple linear model) on Fv/Fm were detected.

### Holobiont oxygen fluxes

Rates of P_gross_, P_net_ and R varied significantly across the experimental conditions (p < 0.001, Kruskal-Wallis test) (**Figure 2A; <u>Supp. Table S2</u>**). Under light conditions, cold-stressed anemone holobionts had negative net photosynthesis while control holobionts were net productive (−0.270 ± 0.279 µmol O₂·mg⁻¹·h⁻¹ compared to controls: 0.159 ± 0.0829 µmol O₂·mg⁻¹·h⁻¹) (**Figure 2A; <u>Supp. Table S2</u>**). Importantly, this decline in P_net_ was not a result of increased dark respiration. Instead, cold stress significantly reduced respiration rates in cold-stressed individuals by 64% (−0.0678 ± 0.0102 µmol O₂·mg⁻¹·h⁻¹) compared to controls (−0.180 ± 0.0750 µmol O₂·mg⁻¹·h⁻¹) (p < 0.001, Conover’s test). Taken together, our results thus indicate a complete loss of P_gross_ in cold-stressed anemone holobionts, which did not show detectable gross O₂ production in the light (0.00 ± 0.00 µmol O₂·mg⁻¹·h⁻¹) compared to controls (0.339 ± 0.145 µmol O₂·mg⁻¹·h⁻¹) (p < 0.001, Conover’s test) (**<u>Supp. Table S2</u>**).

### Superoxide Dismutase (SOD) activity and metabolite profiles of sea anemone holobionts

Superoxide dismutase (SOD) activity doubled in cold-stressed individuals (42.6 ± 16.6 U·mg⁻¹ host tissue protein) compared to controls (20.3 ± 7.65 U·mg⁻¹ host tissue protein) (**Figure 2D; <u>Supp. Table S2</u>**). In addition, we observed a clear effect of cold stress on the overall exometabolite profiles of *A. couchii* holobionts (PERMANOVA; positive ionization mode: F = 21.969, R^2^ = 0.54965, *p* = 0.001; negative ionization mode: F = 29.842, R^2^ = 0.53658, p = 0.001) (**Figure 2B,E; <u>Supp. Table S4</u>, <u>S5</u>**). Together, PC1 and PC2 explained 37.2% and 37.4% of the overall variation in positive and negative ionization datasets, respectively.

OPLS-DA for both ionization modes confirmed the robustness and predictability of the models obtained and allowed us to proceed with biomarker identification (positive ionization mode: R2Y = 0.893, Q2 = 0.803; negative ionization mode: R2Y = 0.893, Q2 = 0.803; permutation test: p < 0.001; **Supp. Figure S5,6**). After manually checking each of the most discriminant VIPs to eliminate potential artifacts, we identified 25 metabolites of interest (VIP score >1.6), most of them exhibiting increased relative abundances in both positive and negative ionization modes, indicating a strong contribution to separation between treatments (**Figures 3, 4; <u>Supp. Table S6a,b</u>; <u>Supp. Table S7</u>**). Metabolite annotation revealed a strong enrichment in dipeptides (18 features), particularly those containing branched-chain amino acids (**Figure 3**). The MS/MS spectra displayed the characteristic leucine immonium ion (m/z 86.097), supporting the presence of a terminal leucine/isoleucine residue (Falick *et al*., 1993). Leucine and isoleucine are isobaric compounds and produce highly similar MS/MS fragmentation spectra, which prevents their unambiguous discrimination under the experimental conditions used (Falick *et al*., 1993). These compounds displayed high fragmentation similarity scores (Tanimoto coefficients 81-98%) and low mass errors (median < 10 ppm; **<u>Supp. Table S6a,b</u>**), supporting confident metabolite annotation. Six of these compounds, all of which were dipeptides, appeared as VIP features in both ionization modes, highlighting their relevance (indicated in bold in **<u>Supp. Table S6a,b</u>**). Additional chemical classes with significantly increased intensities in the cold-stressed holobiont such as fatty acyl carnitines (2 features in positive mode), sphingolipids, specifically ceramides (1 and 4 features in positive and negative modes, respectively), and a glycerophosphocholine (**Figure 4**). In addition, 20 variables were annotated as glycolipids, including glycosyldiacylglycerols (GDGs) and other fatty acyl glycosides (mean±SE log2FC −2.2±0.3 and −1.8±0.1, respectively), all of which exhibited reduced abundances in cold-stressed *A. couchii* holobionts (**<u>Supp. Table</u> <u>S6a,b</u>**; **<u>Supp. Table S7</u>**; **Supp. Table S8**).

**Figure 3.**
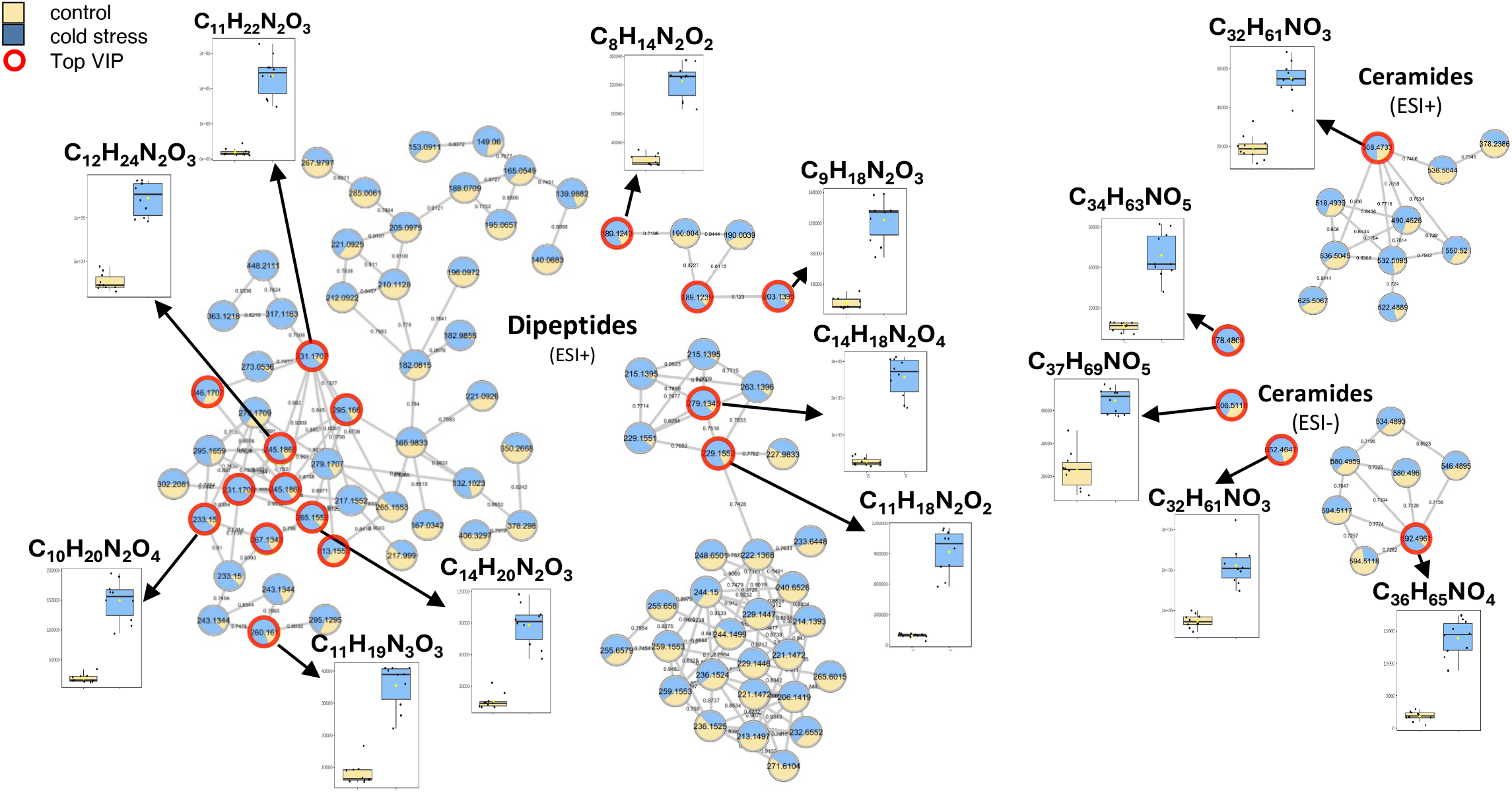
Molecular Network generated from the datasets obtained from positive (UHPLC-ESI(+)-MS/MS and negative ionization (UHPLC-ESI(-)-MS/MS) modes for *A. couchii* holobionts showing dipeptide and ceramide clusters containing VIP features differentially abundant under cold stress. Y-axes of insets denote relative abundances (based on peak areas) of metabolites. ESI+: positive ionization mode; ESI(-): negative ionization mode.

**Figure 4.**
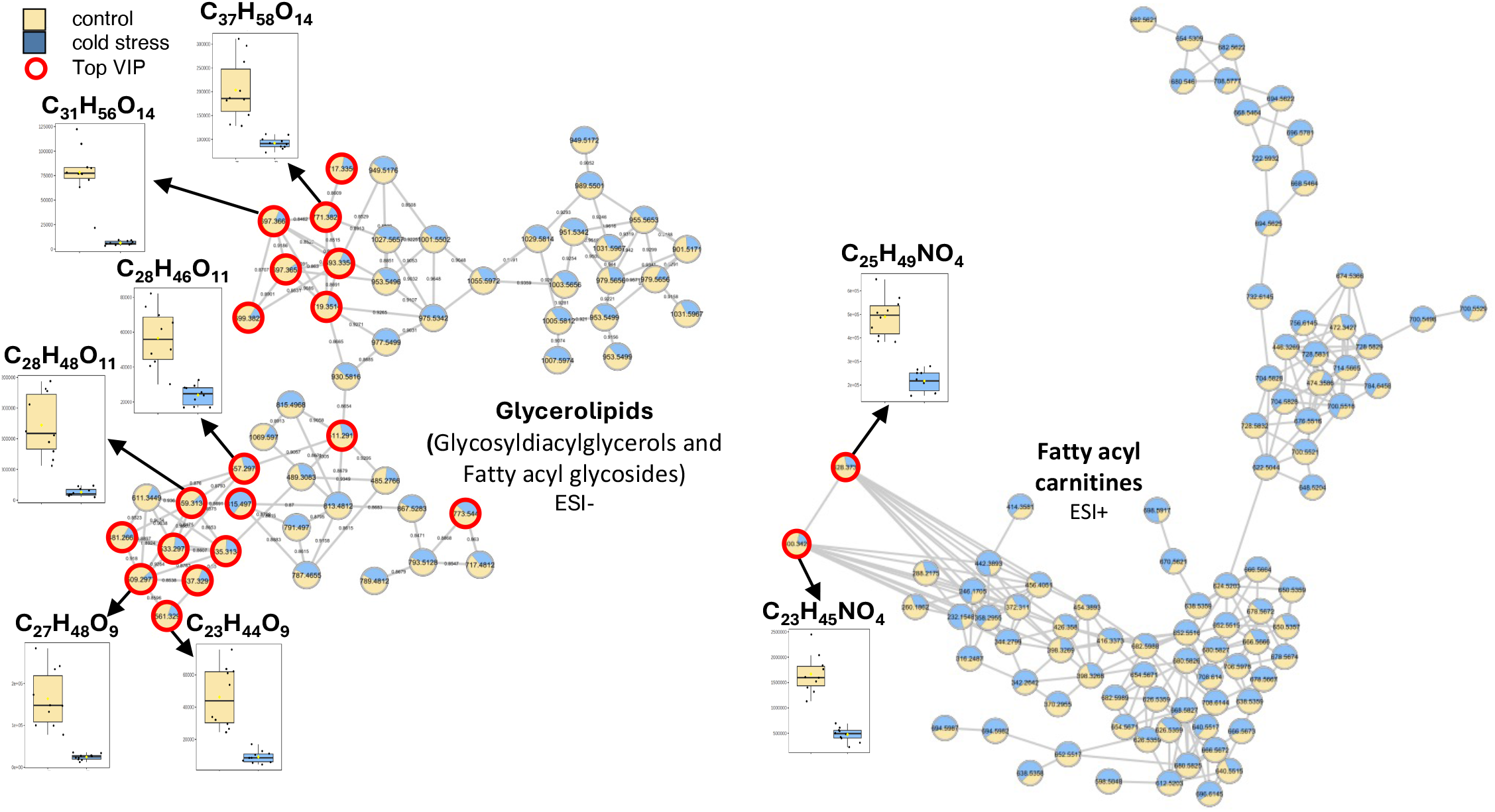
Molecular Network generated from the datasets obtained from positive (UHPLC-ESI(+)-MS/MS and negative ionization (UHPLC-ESI(-)-MS/MS) modes for *A. couchii* showing glycerolipid and fatty acyl carnitine clusters containing VIP features differentially abundant under cold stress. Y axes of insets denote relative abundances (based on peak areas) of metabolites. ESI+: positive ionization mode; ESI(-): negative ionization mode.

## Discussion

The cnidarian-algae symbiosis, a mutualistic relationship between the animal host and the endosymbiotic Symbiodiniaceae, is highly sensitive to a range of environmental fluctuations including nutrients, pollution, salinity, but in particular of temperature (Weis 2008; Davy et al. 2012; Dubé et al. 2025). Due to the impacts of rapidly accelerating anthropogenically-driven climate change, our understanding of high temperature stress on this symbiosis - in particular the stress response known as coral bleaching - is relatively advanced (Hoegh-Guldberg, 1999; Hughes *et al*., 2017; Rädecker *et al*., 2021). Conversely, despite low temperatures being a long-documented cause of bleaching in photosymbiotic Cnidaria (Steen & Muscatine, 1987; Saxby *et al*., 2003; Roth *et al*., 2012), the underlying mechanisms remain largely unknown.

Here we set out to characterize the effects of cold stress on the physiological and metabolic responses of the temperate photosymbiotic *Aiptasia couchii* holobiont. While our results are testimony to the remarkable resistance of these temperate holobionts to temperatures well below their annual average minimum (**Supp. Figure S1B**), they are clearly not immune to the effects of severe cold stress. Indeed, the gradual exposure to increasing cold stress elicited a clear stress phenotype. However, stable Fv/Fm concomitant with reduced endosymbiont population density, loss of photosynthetic pigment content, absence of P_gross_, and altered metabolite profiles paint an intriguing picture of holobiont breakdown. In this scenario, the breakdown of symbiotic nutrient cycling and resulting onset of host starvation closely resemble processes during heat-induced bleaching. Of note, in our study the bleaching was not obvious except for white patches at the tips of the tentacles. The clear reduction of algal symbiont densities and chl *a* content (when normalized to host tissue protein) however are classical hallmarks of cnidarian bleaching (Jones, 1997; Jones & Hoegh-Guldberg, 1998), which was likely masked by the concomitant contraction of animal tissues. Such ‘invisible’ bleaching was previously observed in the heat-stressed photosymbiotic jellyfish *Cassiopea andromeda*, where symbiont loss was obscured by reduced biomass and subsequent shrinkage of the starving host (Toullec *et al*., 2024). In this light, invisible bleaching responses of photosymbiotic Cnidaria (and potentially other photosymbiotic invertebrate systems) which do not form rigid, calcareous skeletons may be common, but largely overlooked. Although not within the scope of this work, these observations prompt the question whether the interpretation of high ecological resilience of such photosymbiotic, ahermatypic organisms largely based on visual bleaching cues are appropriate.

### Cold stress decouples light and dark reactions of photosynthesis in algal symbionts

Bleaching responses to transient cold stress have been reported in a range of photosymbiotic Cnidaria, including the tropical anemone *Aiptasia pulchella (*Muscatine *et al*. 1991; Gates *et al*., 1992) and reef-building corals (Gates *et al*., 1992; Saxby *et al*., 2003; Marangoni *et al*., 2021; El-Khaled *et al*., 2025). Indeed, cold stress is known to induce bleaching in photosymbiotic Cnidaria so reliably that standard protocols to render sea anemone lab models aposymbiotic (endosymbiont-free) typically include an initial cold shock step at 4°C over four hours (Xiang *et al*., 2013). Importantly, despite the striking loss of gross photosynthesis, the remaining symbiont population in cold-stressed *A. couchii* holobionts retained a high photochemical efficiency, i.e., were potentially viable. This specific set of physiological responses of the algal symbionts - the inhibition of photogenic O_2_ production concomitant with an intact PSII - strikingly parallels chilling responses in plants and microalgae following exposure to sublethal cold stress, suggesting a decoupling of light and dark reactions of photosynthesis (Van Hasselt & Van Berlo, 1980). In this scenario, the collapse of photosynthesis is not a result of photodamage to PSII reaction centers but due to a downstream impairment of carbon metabolism due to reduced enzymatic efficiency (Maxwell & Johnson, 2000; Baker, 2008). The resulting activation of cyclic electron flow around PSI sustains the proton motive force and ATP production while limiting linear electron flow and O_2_ evolution (Munekage *et al*., 2004; Baker, 2008). Briefly, the limited electron flux from PSII protects reaction centers from over-reduction and photodamage in the absence of reduced carbon assimilation (Allen, 2003; Kramer *et al*., 2004). Thereby, PSII efficiency is preserved while photosynthetic electron transport and O_2_ evolution are functionally down-regulated due to cold-induced metabolic constraints. While the present study did not assess photobiological responses beyond the maximum quantum yield of PSII and holobiont O_2_ fluxes, certain changes in the metabolite profiles of cold-stressed *A. couchii* holobionts provide indirect support for the decoupling of light and dark reactions of photosynthesis in the algal endosymbiont population being at the root of symbiotic breakdown (**Figure 5**). Lastly, this observation challenges the reliability of the maximum quantum yield as the sole photophysiological response parameter of (cold) stress in photosymbiotic holobionts.

**Figure 5.**
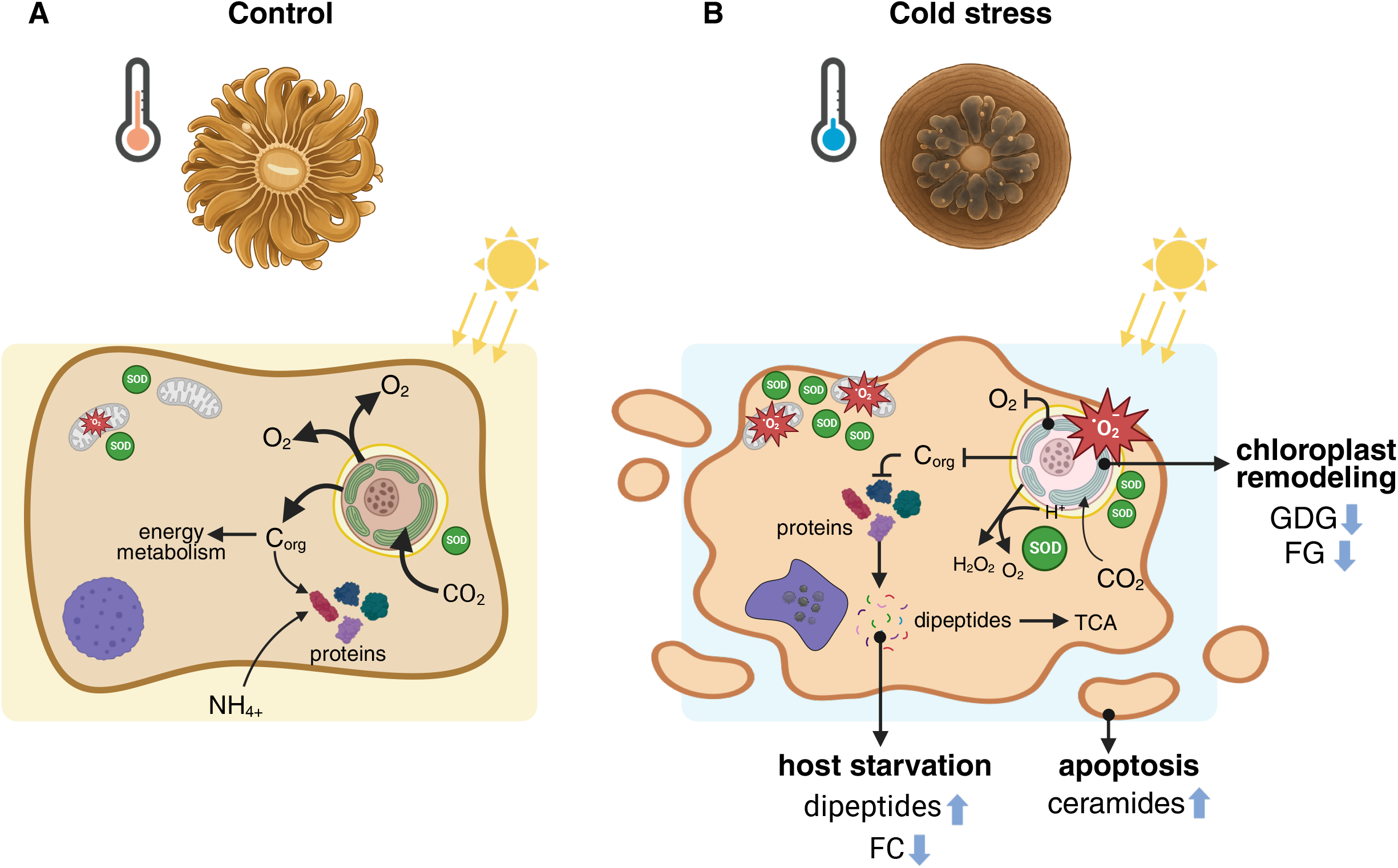
Reconstruction of the chilling-induced responses of the photosymbiotic cnidarian holobiont. Cellular processes and symbiotic nutrient cycling under (**A**) unperturbed conditions and (**B**) cold stress based on physiological and metabolic responses. CO_2_: carbon dioxide; H_2_O_2_: hydrogen peroxide; FG: fatty acyl glycoside; FC: fatty acyl carnitine; GDG: glycosyldiacylglycerol; O_2_: oxygen molecule; •O_2_^-^: superoxide radical; SOD: superoxide dismutase.

### Alteration of thylakoid membranes in cold-stressed algal symbionts

The thylakoid membrane’s lipid composition is essential for cold tolerance in photosynthetic organisms, with glycosyl diacylglycerols (GDGs) being key components (Moellering *et al*., 2010; Li *et al*., 2020). In contrast to the common response of increasing GDGs or lipid unsaturation for cold acclimation in other microalgae and plants (Yi-Bin *et al*., 2017; Zhao *et al*., 2024), our study found significantly reduced relative abundances of GDGs and fatty-acyl glycosides in cold-stressed temperate algal symbionts. Of note, the majority of these metabolites experienced reductions in relative abundances compared to control conditions well above 50 % (**Supp. Table S8**). As such, biomass loss of the Symbiodiniaceae under cold stress alone does not likely explain the observed reductions in GDGs and fatty-acyl glycoside content; rather, the observed reductions in membrane lipids may reflect endosymbiont physiology. Overall, this suggests either an impairment of thylakoid function, thereby reducing cold resistance, or a controlled remobilization of chloroplast lipids to support the repair of damaged photosystem components or generate carbon backbones as alternative energy sources in the absence of photosynthate (Botté *et al*., 2011; Ferrer-Ledo *et al*., 2023). These changes collectively indicate an integrated response where the algal symbiont reconfigures its membrane and carbon partitioning to maintain photosystem stability and cellular economy at reduced enzymatic activity under low temperatures (Willette et al., 2018).

### Chilling-induced disruption of symbiotic nutrient cycling in the *A. couchii* holobiont

A major hallmark of the cnidarian-algae symbiosis is the reciprocal nutrient exchange between the two symbiotic partners: algal endosymbionts transfer a large share of their photosynthate (predominantly sugars, lipids, and sterols) to the host, thereby constituting the major source of energy for the host metabolism (Muscatine 1990; Tremblay *et al*. 2012; Ezzat *et al*. 2015; Yee *et al*. 2025). In turn, CO_2_ from host respiration fuels photosynthesis by the endosymbionts, enabling efficient carbon recycling in the intact symbiosis. Notably, the dynamics of symbiotic nutrient cycling are highly sensitive to environmental change (Muller-Parker & Davy, 2001; Rädecker *et al*., 2021; Toullec *et al*., 2024). Indeed, cold stress caused the effective arrest of gross photosynthesis in photosymbiotic *A. couchii* holobionts in the present study, inevitably resulting in the subsequent collapse of endosymbiont carbon fixation and translocation to the host, ultimately causing severe carbon limitation of the host (Rädecker *et al*., 2021; Toullec *et al*., 2024).

As a likely consequence of such carbon (i.e., energy) limitation, patterns of altered dipeptide abundances, as reflected in increases in 14 different dipeptides, many of which contained branched-chain amino acids, suggest a mobilization of protein stores by the *A. couchii* host in response to reduced energy input (**Figure 5**). Indeed, proteinogenic dipeptides are products of enzymatic protein degradation (Minen *et al*., 2023). Such proteolysis into smaller peptides, and ultimately free amino acids permits the use of carbon backbones from free amino acids for central carbon metabolism (e.g., the TCA cycle) for oxidation and energy production (Chandel, 2021; Torres *et al*., 2023). Similarly, fatty acyl-carnitines (FC) are products of fatty acid esterification to carnitine as part of the carnitine shuttle, a biochemical system central to energy metabolism that enables long-chain fatty acid transport into the mitochondrial matrix, where they are converted back to acyl-CoA and undergo β-oxidation for energy production (Xiang *et al*., 2025). Importantly, FCs are present in animals, plants, and microalgae, although FC content may be orders of magnitude greater in the former (Bourdin *et al*., 2007; Ballesteros-Torres *et al*., 2019). Today, FCs are widely used as mitochondrial biomarkers of fatty acid oxidation which can reflect altered mitochondrial metabolism in diabetes, cancer, and heart failure patients (Vissing *et al*., 2019; McCann *et al*., 2021). In photosymbiotic cnidarian holobionts specifically, abundances of fatty acyl carnitines have been reported to positively correlate with the maximum quantum yield of algal symbionts under hypoxic conditions (Glass & Barott, 2025); however, the exact roles of these fatty acid esters in cnidarian physiology, cell biology and metabolism remain unknown. In the light of arrested gross photosynthesis and subsequent disruptions to symbiotic nutrient cycling, however, the significantly reduced relative abundances of fatty acyl carnitines likely reflects carbon, i.e. energy limitation of the cold-stressed *A. couchii* holobiont, thereby lending further support to the physiological results and observed increased relative abundances of several dipeptides.

As such, metabolic shifts in the anemone holobiont suggest increased rates of catabolic degradation of energy-rich compounds such as fatty acids and proteins by the host, indicating a shift from net anabolism towards net catabolism in the animal host, as previously observed under heat stress (Oakley *et al*., 2016; Rädecker *et al*., 2021; Pei *et al*., 2022; Haydon *et al*., 2023). Similarly, we propose that cold stress subjects the holobiont to a state of imbalanced carbon economy, with the symbiosis being progressively eroded through the rapid depletion of the host’s energy reserves (**Figure 5**). This finding underscores that despite different phenotypic responses at the holobiont level, the underlying metabolic mechanisms of heat and cold stress responses may be conserved across photosymbiotic Cnidaria, and potentially photosymbioses at large.

### Cold stress increases pre-apoptotic markers in the *A. couchii* holobiont

We observed the increased abundance of ceramides in cold-stressed anemone holobionts. These sphingolipids are commonly associated with cold acclimation through increases in membrane fluidity by qualitative changes in membrane composition (Kariotoglou & Mastronicolis, 2001). However, in the context of thermal stress and host starvation, the accumulation of ceramides as stress biomarkers may be the more likely scenario. Indeed, environmental stress including high temperatures can induce the generation of ceramides, which act as second messengers required for apoptosis via SAPK/JNK signalling in animals (Verheij *et al*., 1996; Okazaki *et al*., 1998). Together with other sphingolipids, the pre-apoptotic ceramides are part of the sphingolipid rheostat, which determines cell fate and thereby the cnidarian heat stress response (Kitchen & Weis, 2017). Indeed, apoptosis is one of several mechanisms by which algal symbionts are lost from the cnidarian tissues during bleaching (Lesser & Farrell, 2004; Dunn *et al*., 2004; Richier *et al*., 2006; Weis, 2008). While we did not perform histological confirmation of apoptotic processes, it is widely recognized that apoptosis acts as part of an innate immune response that recognizes algal endosymbionts damaged by oxidative stress and seeks to remove them by programmed host cell death (Weis, 2008). The doubling of SOD activity in tissue homogenates of cold-stressed anemones in the current study lends indirect support to the notion that oxidative stress in the holobiont increased at low temperatures, which may have inflicted damage on both host and symbiont cells, and which may have given rise to the apoptotic cascade in the animal tissues, thereby contributing to the observed stress and bleaching phenotype (**Figure 5**).

## Conclusion

Here we set out to investigate the effects of cold stress on the physiological and metabolic responses of the photosymbiotic sea anemone holobiont *Aiptasia couchii*. Our results reveal clear responses to cold stress, including changes in symbiont density and O_2_ fluxes, host antioxidant activity and holobiont metabolite profiles, while the photosynthetic efficiency of algal endosymbionts remained relatively stable. Thereby, we provide evidence of a symbiotic breakdown in *A. couchii*, characterized by a collapse of the symbiosis driven by the decoupling of light and dark reactions brought on by chilling responses of the algal symbionts. Ultimately, increases in different glycerolipid classes in holobiont metabolite profiles support the notion of symbiotic breakdown resulting in host starvation due to carbon limitation. Our work highlights that despite contrasting physiological responses to heat and cold stress, both stressors destabilize the symbiosis through similar mechanisms deeply rooted in starvation of the animal host, and which inevitably compromise the ecological resilience of photosymbiotic Cnidaria, and in extension the ecosystems they shape, in a rapidly changing world.

## CODE AND DATA AVAILABILITY

Physiology data, data matrices for metabolite profiles, and code are available on zenodo: https://doi.org/10.5281/zenodo.22076964.

Metabolite data are available in private view on the MassIVE server and will be made publicly available upon publication:

Acquisition in *A. couchii* under positive ionization modes, with and without cold thermal stress: doi:10.25345/C5QV3CH6Z [dataset license: CC0 1.0 Universal (CC0 1.0); password for peer review: AcouchiiPos]. Acquisition in *A. couchii* under negative ionization modes, with and without cold thermal stress. [doi:10.25345/C5VH5CX8V] [dataset license: CC0 1.0 Universal (CC0 1.0); password for peer review: AcouchiiNeg].

## AUTHOR CONTRIBUTIONS

NR and CP conceived the study, designed the experiment and obtained funding. ML, TLD, HG and CSM performed the experiment. ML, TLD, HG, CSM, PIHJ and CP performed field work and/or sampling. ML, TLD, HG, DR and NTB processed samples and obtained data. ML, SC, MR, NR, NTB and CP analysed data. ML, NTB and CP curated data. ML, SC, NR, NTB and CP interpreted data. CP wrote the manuscript with contributions from ML, SC, NR, NTB. All authors provided feedback on the final version of the manuscript.

## ACKNOWLEDGEMENTS

We would like to thank F. Morat, M. Brassier, E. Jobet and J. Kotarba for support with administrative and logistic matters. We acknowledge the MSXM/Bio2Mar technical facilities for the mass spectrometry data acquisition. We further thank P. Motta of the Direction Interrégionale de la Mer Méditerranée for the swift issue of sea anemone collection permits, and D. de Beer (Max Planck Institute for Marine Microbiology) and C. Mathiot (PureOcean Fund) for insightful discussions on chilling injury and photophysiology in higher plants, respectively. Finally, the authors would like to express their gratitude to the editor and the anonymous reviewer, whose suggestions helped improve the manuscript. Figure 5 was generated by CP using BioRender Pro licensed to CP (https://www.BioRender.com), artistic renderings of sea anemone phenotypes from photos were generated in ChatGPT.

## CONFLICT OF INTEREST

None declared.

All the research complies with applicable laws on sampling from natural populations and animal experimentation (including the ARRIVE guidelines).

## Ethics Approval Statement

Ethical approval was not required because *Aiptasia couchii* is a non-cephalophod invertebrate not covered by French regulations governing experimental procedures on animals. Wild specimens were collected under DIRM Méditerranée permit 205.

