## Supplementary Material for "Chilling injury to algal symbionts induces host starvation and metabolic reorganization in a temperate cnidarian"

### Supplementary information

Test experiment to determine low temperature stress tolerance. To assess the final temperature for the main cold stress experiment, we performed a four week long experiment during which cold stress was applied through gradual, non-linear downramping of incubator temperatures from 17°C to 5°C (**Supp. Figure S2A**). Phenotyping of holobionts along with the maximum quantum yield (Fv/Fm) of endosymbionts in hospite was assessed weekly. Reductions in Fv/Fm became apparent beginning with exposure to 6°C in week 3 of the test experiment, which persisted until the end of the experiment (**Supp. Figure S3**). However, as sea anemone mortality became apparent shortly after the final temperature ramping to 5°C, we selected 6°C as a final temperature for the main experiment and reduced the duration to 4 weeks.

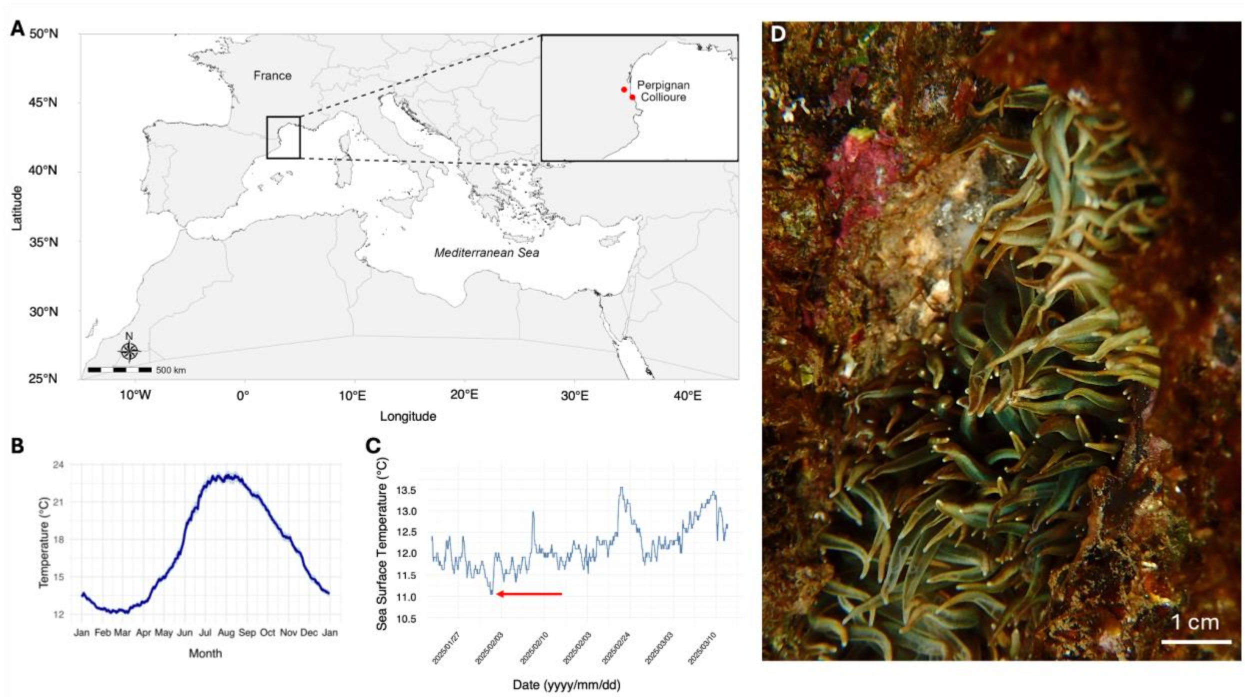


**Supplementary Figure S1. Overview of collection site, *in situ* and experimental temperatures, and experimental organisms. A**. Collection site of sea anemones (*Aiptasia couchii*) off Collioure, France, in the Western Mediterranean Sea. **B**. Mean monthly sea surface temperatures (SST; °C) off Collioure, France, Western Mediterranean Sea (2015 - 2025), obtained from Copernicus Marine Service. Values presented as mean ± SE. **C.** *In situ* sea surface temperatures during the coldest months of the year 2025. The red arrow indicates the lowest temperature during the collection period (01/2025-03/2025). **D**. *In situ* image of *A. couchii* anemones in their natural rocky shore habitat at the collection site (photo by C. Pogoreutz).

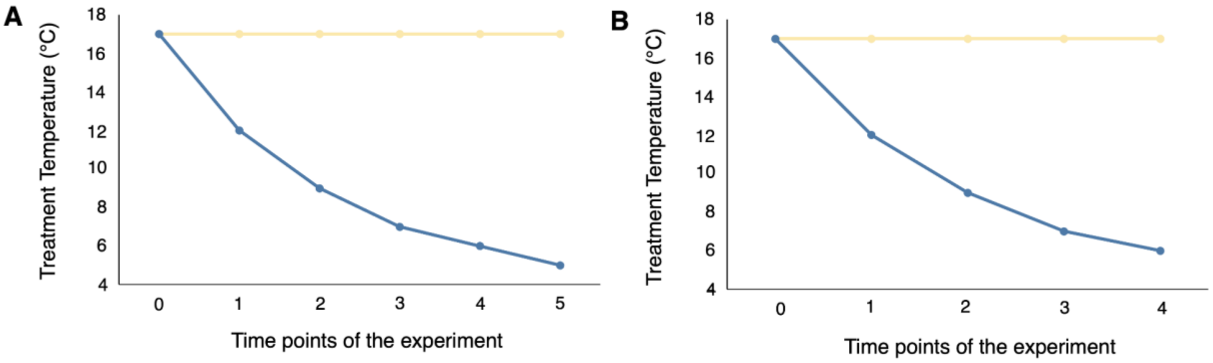


**Supplementary Figure S2.** Progressive, non-linear temperature reduction protocol for the test **(A)** and main **(B)** experiment. Control sea anemones were maintained at 17°C. The cold-stressed anemones in the main experiment experienced a cold stress as follows: 17°C, 12°C, 9°C, 7°C, and 6°C; in the test experiment, the temperature was ramped to a final temperature of 5°C in week 5.

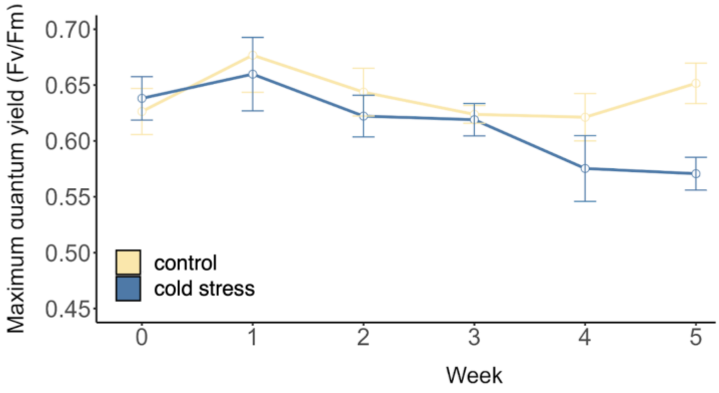


**Supplementary Figure S3. Maximum quantum yield (Fv/Fm)** of algal symbionts *in hospite* observed in test experiment to determine sublethal cold stress temperatures for the main experiment.

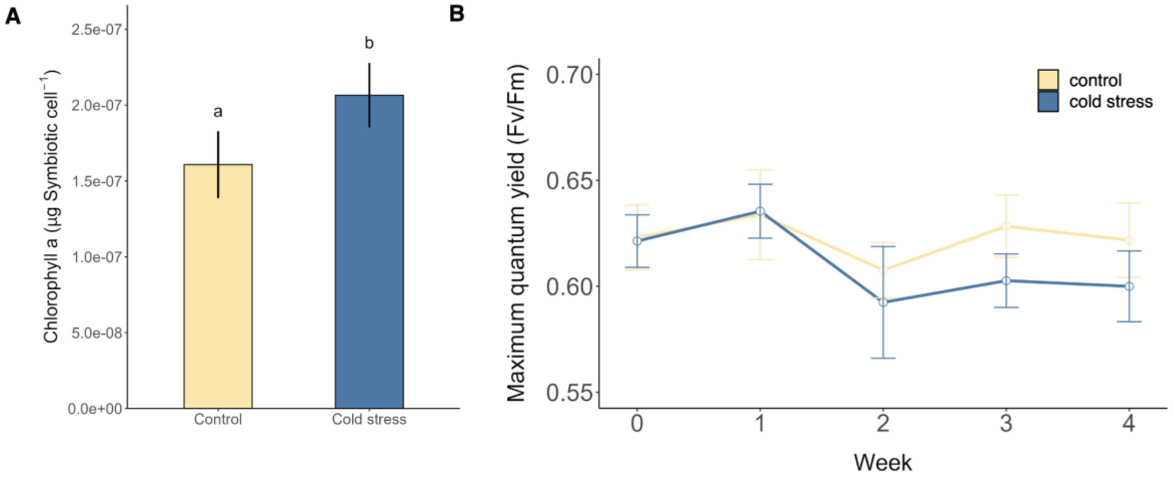


**Supplementary Figure S4. Physiological responses of algal symbiont communities in cold-stressed *A. couchii* holobionts. A**. Chlorophyll *a* content per Symbiodiniaceae cell (µg. symbiont cell$^{-1}$) was significantly increased under cold stress (n = 9 per condition). **B**. No significant effects of cold stress on the Fv/Fm of algal symbionts *in hospite* were observed. Measurements were conducted weekly over four weeks, just prior to the next step in the progressive, non-linear temperature decrease; these steps : 17°C, 12°C, 9°C, 7°C and 6°C. Letters indicate statistically significant differences between groups ($p$ < 0.05). Data are presented as mean ± standard error.

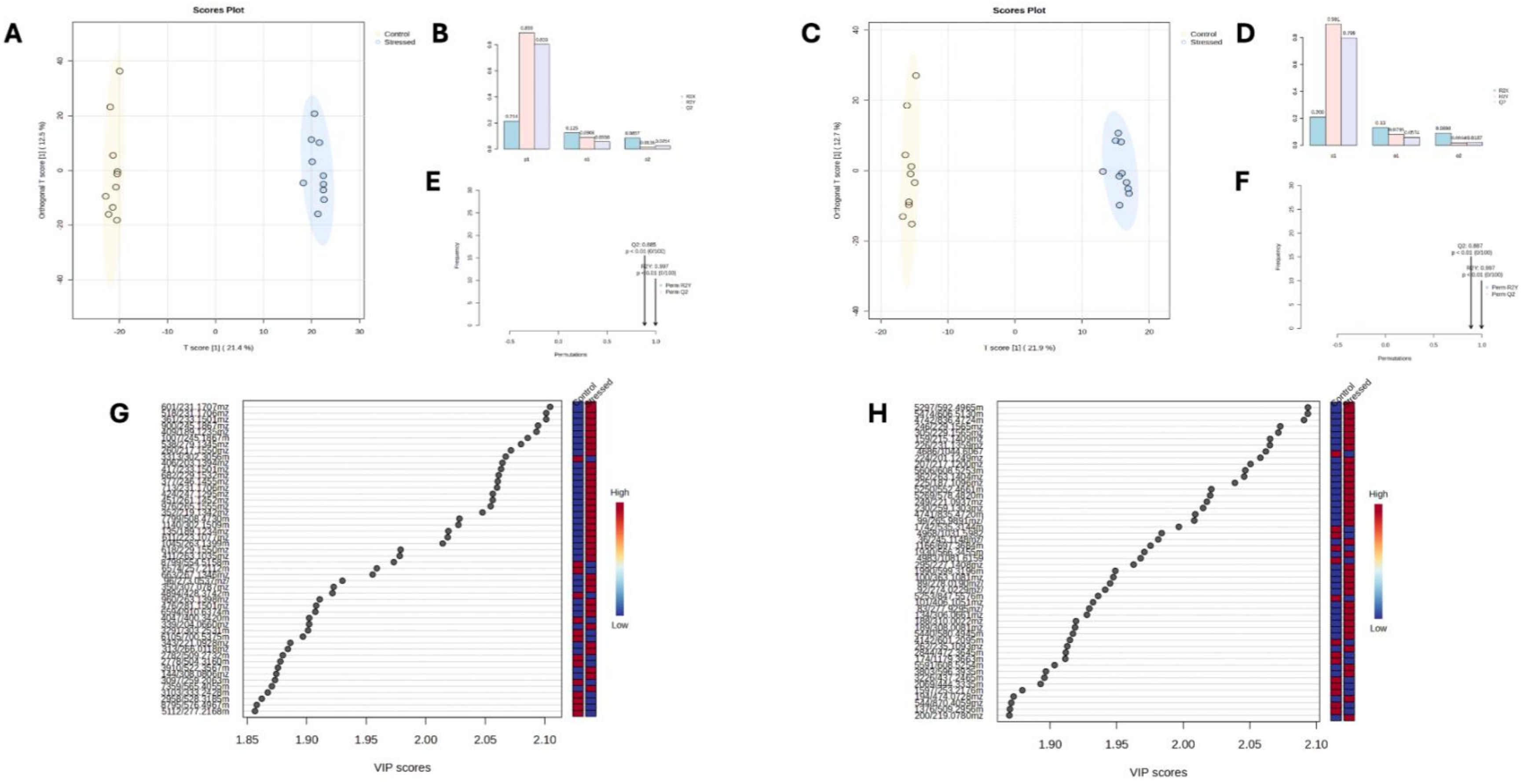


**Supplementary Figure S5.** OPLS-DA score plots and OPLS-DA model validations highlighting discriminant metabolites between control and stressed anemones on positive (A, B, E, G) and negative (C, D, F, H) ionization modes.

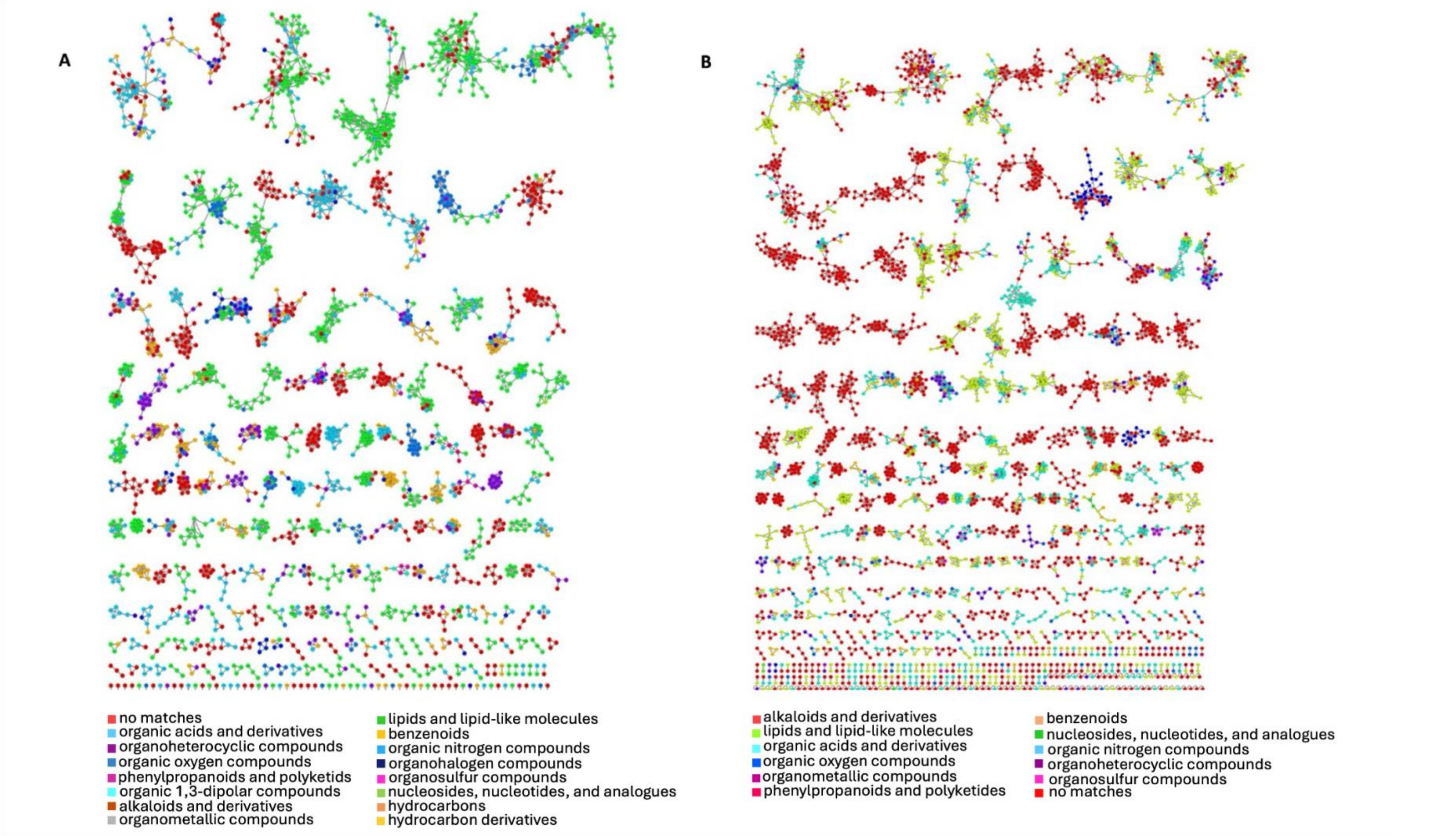


**Supplementary Figure S6.** Feature-Based Molecular Network (FBMN) and Canopus compound class prediction combination in **A.** positive and **B.** negative ionization mode. Nodes are colored based on their chemical superclass.

### Supplementary Tables

**Supplementary Table S1.** Mean annual summer surface temperatures (SST; °C) off Collioure, France, Western Mediterranean Sea (2015 - 2025), obtained from Copernicus Marine Service. Values presented as mean ± SE.

| | **SST (°C)** |
|---|---|
| **2015** | 16.73±1.12 |
| **2016** | 16.72±1.04 |
| **2017** | 16.70±1.06 |
| **2018** | 17.31±1.31 |
| **2019** | 17.57±1.34 |
| **2020** | 16.95±1.19 |
| **2021** | 16.91±1.17 |
| **2022** | 17.55±1.32 |
| **2023** | 17.65±1.27 |
| **2024** | 17.21±1.19 |
| **2025** | 17.43±1.28 |
| **2015-2025** | 17.16±0.40 |

**Supplementary table S8.** Fold changes and log2(FC) obtained for significantly differentially abundant glycosyldiacylglycerols and other fatty acyl glycosides. CCP: Compound class probability; FC: fold change; FG: fatty acyl glycoside; GDG: glycosyldiacylglycerols; VIP: Variable Importance in Projection.

| Code name (MZmine) | Test | FC | log2(FC) | raw.pval | FDR-adjust. pval | Compound classes | CCP | VIP Score |
|---|---|---|---|---|---|---|---|---|
| 1166/697.3684mz/10.32min | OPLS-DA | 0.086262 | -3.5351 | 5.9148e-08 | 4.9148e-07 | GDG | 0.840 | 1.97 |
| 1550/559.3144mz/11.01min | OPLS-DA | 0.11272 | -3.1492 | 2.938e-06 | 1.1673e-05 | GDG | 0.908 | 2.42 |
| 1376/509.2956mz/10.73min | OPLS-DA | 0.16522 | -2.5975 | 1.5536e-06 | 8.7440e-06 | FG | 0.810 | 1.87 |
| 1905/561.3293mz/11.49min | OPLS-DA | 0.21209 | -2.2372 | 1.4974e-05 | 2.9948e-05 | FG | 0.992 | 1.74 |
| 908/481.2660mz/9.64min | Volcano | 0.21409 | -2.2237 | 0.00085378 | 0.00085378 | FG | 0.987 | |
| 832/833.4045mz/9.38min | Volcano | 0.21642 | -2.2081 | 0.00049158 | 0.00085378 | GDG | 0.956 | |
| 2088/611.3458mz/11.79min | Volcano | 0.2255 | -2.1488 | 8.6188e-05 | 0.00011492 | GDG | 0.966 | |
| 2401/563.3464mz/12.17min | Volcano | 0.23939 | -2.0626 | 6.0549e-05 | 8.6499e-05 | FG | 0.955 | |
| 1297/771.3848mz/10.60min | OPLS-DA | 0.24006 | -2.0585 | 0.00058593 | 0.00065103 | GDG | 0.749 | 1.72 |
| 1084/697.3686mz/10.14min | Volcano | 0.25311 | -1.9822 | 2.7771e-05 | 4.3372e-05 | GDG | 0.840 | |
| 2138/537.3287mz/11.85min | Volcano | 0.25347 | -1.9801 | 2.7964e-05 | 4.3372e-05 | FG | 0.950 | |

| | | | | | | | | |
|---|---|---|---|---|---|---|---|---|
| 1929/563.3227mz/11.52min | Volcano | 0.25527 | -1.9699 | 0.00014019 | 4.3372e-05 | FG | 0.955 | |
| 1742/535.3144mz/11.25min | OPLS-DA | 0.28294 | -1.8214 | 2.4835e-08 | 4.3372e-05 | FG | 0.640 | 1.77 |
| 866/717.3338mz/9.49min | OPLS-DA | 0.31456 | -1.6686 | 0.00077484 | 4.9670e-07 | FG | 0.686 | 1.65 |
| 1041/531.2834mz/10.04min | OPLS-DA | 0.3272 | -1.6117 | 3.5018e-06 | 1.1673e-05 | GDG | 0.720 | 1.84 |
| 1295/533.2992mz/10.60min | OPLS-DA | 0.33428 | -1.5809 | 2.8192e-05 | 1.1673e-05 | FG | 0.842 | 1.72 |
| 1604/535.3141mz/11.09min | Volcano | 0.35615 | -1.4894 | 1.0886e-05 | 4.3372e-05 | FG | 0.518 | |
| 1247/511.2910mz/10.50min | OPLS-DA | 0.44863 | -1.1564 | 1.7488e-06 | 2.7215e-05 | FG | 0.559 | 1.85 |
| 1244/557.2959mz/10.50min | OPLS-DA | 0.4593 | -1.1225 | 6.3515e-06 | 8.7440e-06 | GDG | 0.941 | 1.78 |
| 1149/509.2765mz/10.29min | OPLS-DA | 0.47408 | -1.0768 | 1.3437e-05 | 1.8147e-05 | FG | 0.571 | 1.75 |

**Supplementary Tables S2-S7** are publicly available on https://doi.org/10.5281/zenodo.22076964

**Supplementary Table S2.** Physiological response parameters of *Aiptasia couchii* holobionts under control and cold-stressed conditions on day 28 of the experiment. Symbiont_count: Endosymbiont cell densities normalized to host tissue protein content; chla_protein: chlorophyll *a* content of endosymbionts normalized to host tissue protein content; chla_cell: chlorophyll *a* content of endosymbionts normalized to endosymbiont cell numbers: O2_gross: gross photosynthesis rates of *A. couchii* holobionts; O2_net: net photosynthesis rates of *A. couchii* holobionts; O2_resp: respiration rates of *A. couchii* holobionts. SOD: SOD activity in *A. couchii* tissue homogenates.

**Supplementary Table S3.** Weekly maximum quantum yields (Fv/Fm) of dark-adapted *Aiptasia couchii* holobionts under control and cold-stressed conditions. Temperatures in cold-stress treatment: Weeks 0-4 = 17, 12, 9, 7 and 6 °C, respectively. Fv.Fm: Fv/Fm values.

**Supplementary Table S4.** Absolute abundance data (peak area) for metabolite features identified in the positive ionization mode.

**Supplementary Table S5.** Absolute abundance data (peak area) for metabolite features identified in the negative ionization mode.

**Supplementary Table S6.** Annotation table for 25 significantly and most differentially regulated VIP metabolites for positive and negative ionization mode, respectively.

**Supplementary Table S7.** Output from Wilcoxon Rank Sum tests to assess statistical differences in the intensity of statistically significantly differentially regulated VIP features between control and cold-stressed sea anemone holobionts.
